# A New Method for Primary Culture of Mouse Dorsal Root Ganglion Neurons

**DOI:** 10.1101/466110

**Authors:** Tiantian Dong, Shigang Li, Wei Liu, Mengzhen Yan, Jie Yu, Xiaozhuo Deng

## Abstract

In order to establish a simple and highly purified method for primary culture of mouse dorsal root ganglion neurons(DRGn), in this study, the DRGn of young mice were obtained by collagenase type I and trypsin digestion. Then the DRGn were obtained by immunocytochemical staining of mouse neuron specific enolase (NSE) monoclonal antibody, while using flow cytometry to further detect the positive rates of DRGn. The cultured primary DRGn grew well and had a purity of about 83.72%. The DRGn survival time was 60 days when cultured in DMEM medium containing nerve growth factor (NGF). The culture scheme is simple and stable, and a large number of high purity DRGn could be cultured, which provides a reliable model for further study of nerve cells.

## Introduction

Spinal DRGn are afferent neurons of peripheral nervous system and primary neurons of sensory conduction system. With the increasing research on the nervous system, such as the pain mechanism of diseases[1-3], spinal cord injury repair[4,5], electrophysiological research[6-9], it is necessary to establish a simple and easy-to-obtain primary nerve cell culture scheme. Based on the previous studies [10-13], this scheme simplifies the cultivation process, aiming to cultivate a large number of high-purity nerve cells and providing a stable and convenient model for neuropathophysiology research.

## Methods and procedures

### Animals

The healthy male C57BL/6 mice aged 6~8 weeks were selected from the animal center of China Three Gorges University. All procedures in the animal studies were approved by the Animal Care and Use Committee at China Three Gorges University.

### Isolation and acquisition of DRGn

C57BL/6 mice were sacrificed by cervical dislocation and soaked in 75% alcohol for 3 minutes. The mice were dissected from the back in order to take out the spinal column and were put in the plate of 4°C Hank’s solution. The spinal columns were divided into two halves by ophthalmic scissors along the midline of the spine. The spinal cords were gently moved by microforceps to make the DRG exposure. Each DRG was collected in a centrifuge tube while the nerve roots were cut off. An appropriate amount of 0.125% collagenase type I was added into centrifuge tube to digest DRG and was shaken at 37°C for 30-50 minutes, and then it was centrifuged to remove collagenase type I. Next, the previous step was repeated before the collagenase type I was removed again. Finally, a proper amount of 0.25% trypsin was added into the centrifuge tube before it was shaken for about 20-30 minutes at 37°C.

After digestion, the supernatant was removed via centrifugation and was added to DMEM complete medium: 90% DMEM (Gibco), 10% horse serum (Gibco, 26050-088), 1% penicillin/streptomycin (Hyclone) and 100μg/L NGF (novoprotein, CK21). Repeatedly blew DRGn with a sterile tiny glass pipette to form a single cell suspension, and then inoculated DRGn into the culture dish. The culture dishes were coated with 0.1mg/ml Poly-D-lysine solution (Sigma, 27964-99-4) one day earlier, and DRGn were cultured in 37°C, 5% CO^2^ incubator.

After 24 hours of culture, the medium in the culture dishes was change entirety, and the medium was changed every 3 days. The DRGn at different time stages were observed under an Olympus microscope and their morphological changes were also recorded.

### Immunocytochemistry

After 7 days of culture, the DRGn were washed for 3 times with phosphate-buffered saline(PBS), 0.01M, pH7.4, and were fixed with 4% paraformaldehyde at room temperature for 40 minutes, then were washed for 3 times with PBS for 5 minutes each time. 0.1% Triton solution was submerged in for 30 minutes at room temperature, and was washed for 3 times with PBS for 5 minutes each time. To block non-specific protein, 10% goat serum was submerged in for 30 minutes at room temperature before it was then removed. The DRGn were incubated overnight at 4°C with NSE monoclonal antibody (Abcam, ab16808) diluted at the ratio of 1:100. The primary antibody was recovered and washed for 3 times with PBS, and was incubated with goat anti-mouse IgG (KERUI, 125435, China) at the ratio of 1:500 for 1 hour at room temperature. The secondary antibody was also recovered and washed for 3 times with PBS, and DAB staining solution was submerged in to stain for 5 minutes at room temperature. After thorough rinse, hematoxylin solution was counterstained for 3 minutes, and then rinsed clean with distilled water. Finally, stained samples were sealed with neutral gum, and were examined with microscope.

### Flow cytometry

7th day DRGn were cultured for flow cytometry. DRGn were collected in eppendorf tubes by digesting and centrifuging. The DRGn were fixed with 4% paraformaldehyde for 10 minutes at room temperature, and paraformaldehyde was removed via centrifugation. The DRGn were incubated in 10% goat serum to block non-specific protein. Next, centrifuged and removed goat serum, and incubated the DRGn with NSE monoclonal antibody (dilution of 1:1000) for 30 minutes at room temperature. The fluorescent secondary antibody goat anti-mouse IgG (Bioss, bs-0296G-AF488, China) was used to incubate DRGn at room temperature in the dark for 30 minutes. For DRGn in the control group, just added 500ul PBS to suspend cells. Flow cytometry 488 laser channels were used to detect.

## Results

### Morphological observation of neurons in DRGn

The primary cultured DRGn adhered slowly and most of them adhered to the wall after 24 h. The DRGn presented a clear spherical shape in the first day. When cultured to the 5th day, most of the DRGn protruded synapses and the characteristic morphology of neurons was obvious. On the 7th day, the DRGn grew vigorously in good condition. The DRGn intertwined into a network with axons and dendrites, and had larger cell bodies, which were round or polygonal. After 15 days, the bodies of DRGn gradually shrank and became irregular, who showed an aging trend, and the cross-links among cells were disordered. The medium was changed semi-quantitatively every 3 days until a plenty of DRGn were still visible on the 60th day (Figure.1).

**Figure 1.**
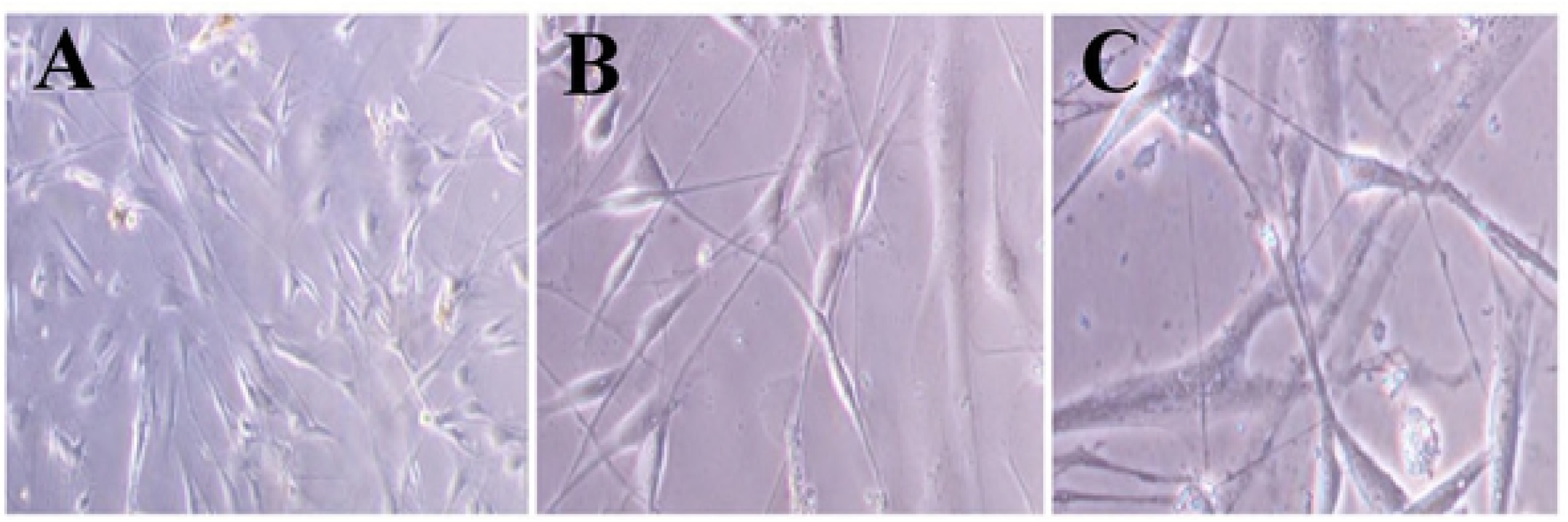
The morphological structure of DRGn were observed under inverted phase contrast microscope on the 7th day of culture:(A) 100Xs (B) 200X;(C) 400X

### Immunocytochemistry of DRGn

The nuclei of all DRGn were stained blue by hematoxylin under an inverted phase contrast microscope (Figure.2). The NSE antibody-labeled DRGn cytoplasms were stained brown. The axons and dendrites of DRGn intersected together and had clear structures. Thus, it could be seen that most of the cells were positive DRGn.

**Figure 2.**
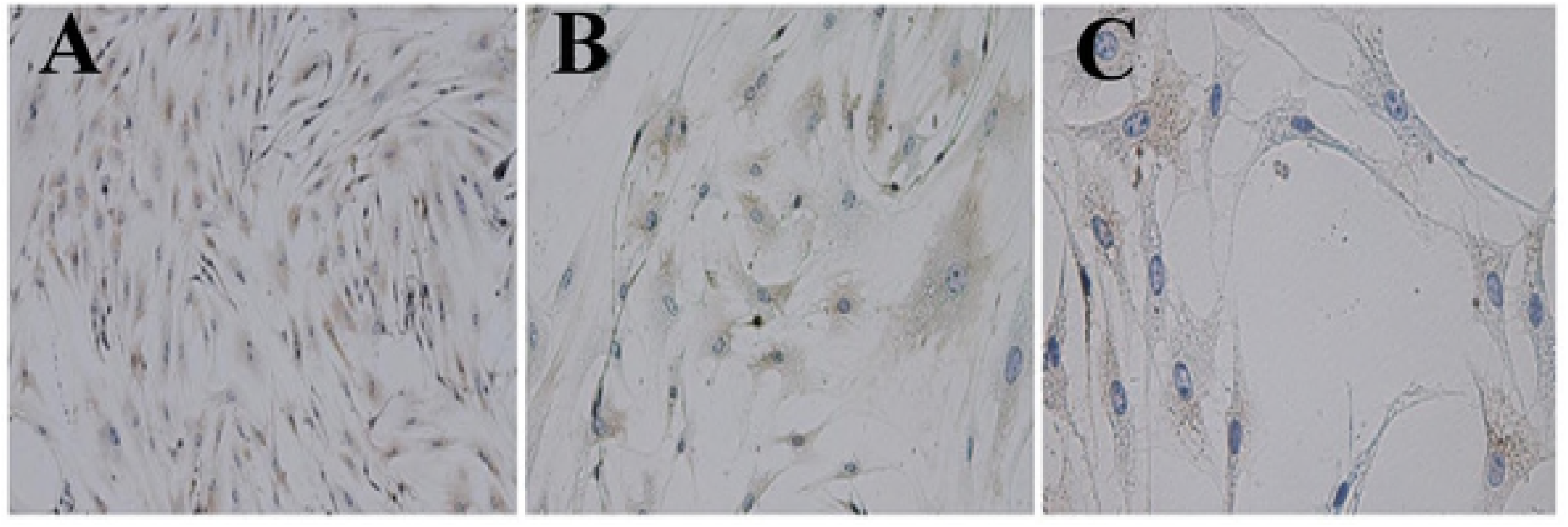
Immunocytochcmical staining showed that the cytoplasm of NSE-labcllcd DRGn were brown and the nuclei stained with hematoxylin were blue: (A) 100X;(B) 200X;(C) 400X

### Purity determination of DRGn

DRGn labeled with NSE monoclonal antibody had high average fluorescence intensity in the experimental group while there was low fluorescence value in the control group with no antibody labeling (Figure.3). The positive rate of DRGn was 83.72% after flow cytometry detection.

## Discussion

The DRGn of the spinal cord are often used to study the mechanism of pain. At present, most of the neurons are cultured from embryonic rats or newborn rats because the infant neurons are easier to survive than the nerve cells of adult rats. However, it is difficult to obtain the infant neurons, because it requires higher experimental conditions and techniques. Therefore, in this experiment, a new DRGn extraction method of young mice was established to solve the above questions. The experimental method was simple and stable, and a large number of DRGn were obtained.

The DRG of young mice was extracted and digested by collagenase type I and trypsin to form single cell suspension for culture. Fibroblasts were the common impurity cells when cultured DRGn. Differential adherence method is commonly used to purify DRGn, meanwhile some DRGn will be lost in this process. Another purification method is to add 5-fluoro-2-deoxyuridine nucleotide and uridine after 24 hours of cultivation to clear the proliferative non-neuronal cells [14, 15]. Although it exerts a certain effect, it also causes partial damage to DRGn. This program, however, has undergone a large number of experiments so as to improve the methods of extracting the primary DRGn of the predecessors. The biggest improvement in this program was that the digestion time was prolonged. The digestion time was 60-90 minutes with collagenase type I twice, and the digestion time of trypsin was 20-30 minutes. The purity of the primary DRGn extracted by this program can reach more than 80%, without the need to add purified drugs. It is concluded that the enzymatic method removes some impurity cells to a certain extent.

10% horse serum and 100μg/L NGF were added to DMEM medium for culturing DRGn. Fetal bovine serum contained mitotic factors, which were often used as serum for culturing proliferating cells. But horse serum can avoid this defect. Therefore, if horse serum were used in nerve cell culture, the growth of fibroblasts could be inhibited to some extent. NGF can maintain the survival of nerve cells, and promote the growth and elongation of axons and dendrites as well, so it is also an essential component in the medium [16-18]. With regard to the selection of Petri dishes, it is advisable to use small Petri dishes, such as 35 mm Petri dishes, because the growth of nerve cells requires certain density, and the cross-linking between axons and dendrites is conducive to the transmission of information among nerve cells.

NSE is a glycolytic enzyme existing in nerve cells and neuroendocrine tissues. It is often used as a marker to identify primary nerve cells [19, 20]. It mainly exists in cytoplasm. Immunochemical staining showed that the cell bodies of the positive neurons were brown, and the positive rate was 83.72% by flow cytometry detection.

The programme of this experiment is easy to operate and obtain materials, which are adult mice instead of embryonic mice. This programme is also convenient to culture DRGn, and can reduce other purification steps. The survival time of DRGn is long, and most of the neurons can still be seen on the 60th day of culture. In conclusion, this experimental method can provide DRGn from adult mice, and provide a new model for the further study of neurology.

## Acknowledgements

This work was supported by National Natural Science Foundation of China (No.81673665, No.81274166). We would like to acknowledge Professor Hebin Tang for his experimental guidance.

## References

1. Shi H, Luo X. 7, 8, 3′-Trihydroxyflavone Promotes Neurite Outgrowth and Protects Against Bupivacaine-Induced Neurotoxicity in Mouse Dorsal Root Ganglion Neurons. Med Sci Monit 2016; 22:2301–2308

2. Yu H, Fischer G, Jia G, et al. Lentiviral gene transfer into the dorsal root ganglion of adult rats. MOL PAIN 2011; 7:63

3. Yu Y, Huang X, Di Y, Qu L, Fan N. Effect of CXCL12/CXCR4 signaling on neuropathic pain after chronic compression of dorsal root ganglion. Sci Rep 2017; 7:5707

4. Wu Z, Li L, Xie F, et al. Activation of KCNQ Channels Suppresses Spontaneous Activity in Dorsal Root Ganglion Neurons and Reduces Chronic Pain after Spinal Cord Injury. J Neurotrauma 2017; 34:1260–1270

5. Zhang J, Liang L, Miao X, et al. Contribution of the Suppressor of Variegation 3-9 Homolog 1 in Dorsal Root Ganglia and Spinal Cord Dorsal Horn to Nerve Injury-induced Nociceptive Hypersensitivity. ANESTHESIOLOGY 2016; 125:765–778

6. Won YJ, Ono F, Ikeda SR. Characterization of Na+ and Ca2+ channels in zebrafish dorsal root ganglion neurons. PLOS ONE 2012; 7:e42602

7. Zhuo M, Gorgun MF, Englander EW. Augmentation of glycolytic metabolism by meclizine is indispensable for protection of dorsal root ganglion neurons from hypoxia-induced mitochondrial compromise. Free Radic Biol Med 2016; 99:20–31

8. Stotz SC, Vriens J, Martyn D, Clardy J, Clapham DE. Citral sensing by Transient [corrected] receptor potential channels in dorsal root ganglion neurons. PLOS ONE 2008; 3:e2082

9. Martinez-Espinosa PL, Wu J, Yang C, et al. Knockout of Slo2.2 enhances itch, abolishes KNa current, and increases action potential firing frequency in DRG neurons. ELIFE 2015; 4

10. Miyano K, Tang HB, Nakamura Y, et al. Paclitaxel and vinorelbine, evoked the release of substance P from cultured rat dorsal root ganglion cells through different PKC isoform-sensitive ion channels. NEUROPHARMACOLOGY 2009; 57:25–32

11. Tang HB, Li YS, Miyano K, Nakata Y. Phosphorylation of TRPV1 by neurokinin-1 receptor agonist exaggerates the capsaicin-mediated substance P release from cultured rat dorsal root ganglion neurons. NEUROPHARMACOLOGY 2008; 55:1405–1411

12. Wu D, Klaw MC, Kholodilov N, et al. Expressing Constitutively Active Rheb in Adult Dorsal Root Ganglion Neurons Enhances the Integration of Sensory Axons that Regenerate Across a Chondroitinase-Treated Dorsal Root Entry Zone Following Dorsal Root Crush. FRONT MOL NEUROSCI 2016; 9:49

13. Jang IJ, Davies AJ, Akimoto N, et al. Acute inflammation reveals GABAA receptor-mediated nociception in mouse dorsal root ganglion neurons via PGE2 receptor 4 signaling. Physiol Rep 2017; 5

14. Kao DJ, Li AH, Chen JC, et al. CC chemokine ligand 2 upregulates the current density and expression of TRPV1 channels and Nav1.8 sodium channels in dorsal root ganglion neurons. J Neuroinflammation 2012; 9:189

15. Ding R, Sun B, Liu Z, et al. Advanced Oxidative Protein Products Cause Pain Hypersensitivity in Rats by Inducing Dorsal Root Ganglion Neurons Apoptosis via NADPH Oxidase 4/c-Jun N-terminal Kinase Pathways. FRONT MOL NEUROSCI 2017; 10:195

16. Cohen MR, Johnson WM, Pilat JM, et al. Nerve Growth Factor Regulates Transient Receptor Potential Vanilloid 2 via Extracellular Signal-Regulated Kinase Signaling To Enhance Neurite Outgrowth in Developing Neurons. MOL CELL BIOL 2015; 35:4238–4252

17. Aloe L, Rocco ML, Balzamino BO, Micera A. Nerve Growth Factor: A Focus on Neuroscience and Therapy. CURR NEUROPHARMACOL 2015; 13:294–303

18. Skaper SD. Nerve growth factor: a neuroimmune crosstalk mediator for all seasons. IMMUNOLOGY 2017; 151:1–15

19. Bharosay A, Bharosay VV, Varma M, et al. Correlation of Brain Biomarker Neuron Specific Enolase (NSE) with Degree of Disability and Neurological Worsening in Cerebrovascular Stroke. Indian J Clin Biochem 2012; 27:186–190.

20. Haque A, Polcyn R, Matzelle D, Banik NL. New Insights into the Role of Neuron-Specific Enolase in Neuro-Inflammation, Neurodegeneration, and Neuroprotection. Brain Sci 2018; 8

